# Isolation of Zika Virus Replication Complex Reveals a Proviral Nuclear Factor

**DOI:** 10.64898/2026.07.06.736844

**Authors:** Peixi Chang, Bhargava Teja Sallapalli, Yanjin Zhang

## Abstract

Zika virus (ZIKV) is an arthropod-borne flavivirus of international public health impact. ZIKV has a positive-sense, single-stranded RNA genome and remodels intracellular membranes to form replication complexes (RCs). The objective of this study was to isolate and characterize the RCs from ZIKV-infected cells and to identify host-cell components recruited to participate in viral replication. Here, we isolated the RCs from ZIKV-infected Vero cells by detergent treatment and flotation centrifugation. Fractional flotation analysis demonstrated that ZIKV proteins NS2B, NS3, and NS5, and ZIKV RNA were present in the detergent-resistant membranous fraction. In contrast, the ER-resident protein calnexin and a mitochondrial protein were present in the detergent-soluble fractions. The isolated RCs were functional for ZIKV RNA synthesis, as shown by quantitative PCR. To determine the components of the RCs, we conducted mass spectrometry analysis and identified numerous cellular proteins. Among them is the replication factor C subunit 2 (RFC2), an accessory protein of DNA polymerase. RFC2 is involved in ATP binding and hydrolysis and may promote cell survival. ZIKV infection increased the RFC2 protein level and induced its relocation to the cytoplasm. RNAi-mediated silencing of RFC2 reduced ZIKV replication. Together, our results provide insights into ZIKV replication and virus-cell interaction.

**IMPORTANCE:** By isolating and characterizing the ZIKV replication complexes (RCs) from infected Vero cells, this research provides valuable insights into the virus’s strategy for hijacking host intracellular membranes and proteins. The identification of ZIKV proteins (NS2B, NS3, and NS5) and RNA in detergent-resistant, lipid-rich membranous structures and the confirmation of functional RCs in RNA synthesis demonstrate successful RC purification. Mass spectrometry and proteomic analysis of the RCs identified numerous cellular proteins that may contribute to ZIKV RC formation and RNA synthesis. Notably, among them is the replication factor C subunit 2 (RFC2), an accessory protein of DNA polymerase in the nucleus. Our results demonstrate that ZIKV increases the level of RFC2 and relocates it to the cytoplasm and that RFC2 plays a role in ZIKV replication, as its depletion reduces viral yield. These results provide insights into ZIKV-cell interactions and potentially facilitate future development of novel antiviral strategies.

## INTRODUCTION

Zika virus (ZIKV) is a reemerging flavivirus primarily transmitted by infected mosquitoes. After causing an epidemic in Central and South America in 2015-2016, ZIKV continues to circulate in tropical regions, necessitating sustained vigilance and preparedness to detect and respond to new cases (1). ZIKV poses a significant public health concern due to its association with severe neurological conditions, such as microcephaly in infants and Guillain-Barré syndrome in adults (2, 3). ZIKV is a small, spherical, enveloped virus containing a positive-sense, single-stranded RNA genome of approximately 10.8 kb (4). Once the virus enters the host cell, its RNA genome is released into the cytoplasm and promptly translated, yielding a polyprotein, which is cleaved into three structural proteins: capsid, precursor membrane (prM), and envelope (E), and seven non-structural proteins (NS): NS1, NS2A, NS2B, NS3, NS4A, NS4B, and NS5 (4).

Like many other positive-sense RNA viruses, ZIKV induces the formation of the membranous replication complexes (RCs), which are generated through invagination of the endoplasmic reticulum (ER) membrane (5–7). The RCs allow spatiotemporal coordination of the different steps of the viral life cycle and protect viral RNA from the cellular antiviral surveillance (8–10). The ZIKV NSPs are involved in ZIKV replication (4, 7). The ZIKV protein NS1 is essential for reorganizing the ER by inserting its hydrophobic regions into the ER membrane to form an RC-like structure, thereby facilitating viral replication (11). As transmembrane proteins, ZIKV NS4A and NS4B serve as scaffolds for the assembly of the replication complex. NS2B is an essential cofactor for the NS3 protease domain. NS3 also has a helicase function, which unwinds the double-stranded RNA (dsRNA) intermediate to facilitate RNA synthesis (12). NS5 functions as the RNA-dependent RNA polymerase (RdRp), synthesizing new viral RNA (13).

Ultrastructural studies showed that ZIKV induces a drastic reorganization of intermediate filament and microtubule networks and reorganizes the ER to form the RCs containing vesicle packets (VPs), zippered ER, and convoluted membranes (CMs), which are thought to increase the local concentration of viral and cellular factors for efficient viral replication (5, 8, 14). Transmission electron microscopy (TEM), combined with 3D reconstruction, provides a 3D overview of the RCs in ZIKV-infected cells (5, 15). However, biochemical studies are required to elucidate RNA synthesis and the molecular functions of host factors in RCs, and such studies necessitate the isolation and purification of functional complexes.

The objective of this study was to establish a purification method for ZIKV RCs and to characterize their components and functions. First, we tested whether the RCs in the cell lysate were resistant to a nonionic detergent. We then isolated the ZIKV RCs from ZIKV-infected cells after detergent treatment by sucrose gradient flotation and further purified them with iodixanol gradient centrifugation. We also performed proteomic analysis of the ZIKV RCs to identify enriched host factors. Among them is the replication factor C subunit 2 (RFC2), an accessory protein of DNA polymerase in the nucleus. In addition, we determined the effects of ZIKV on RFC2 levels and subcellular localization and assessed the effects of RFC2 depletion on ZIKV replication. This study provides insights into ZIKV recruitment of host factors for efficient replication and facilitates the potential development of novel antiviral strategies.

## RESULTS

### The detergent-resistant fraction of ZIKV-infected cells contains RCs

We performed an experiment to determine whether the detergent-resistant fraction contains ZIKV proteins and negative-sense RNA that may be present in RCs (Fig. 1A) (16). Briefly, Vero cells were infected with the ZIKV PRVABC59 strain (GenBank accession number KX377337) (17) at a multiplicity of infection (MOI) of 15 and harvested 24 h post-infection (hpi). The cells were ruptured with a needle and syringe after incubation in a hypotonic buffer, and the nuclei were then removed by centrifugation. The post-nuclear supernatant (PNS), containing membranous structures, was treated with nonionic detergent Triton X-100 or IGEPAL CA-630 on ice and subjected to centrifugation to separate the detergent-soluble and detergent-resistant fractions. Western blotting (WB) results showed that, like soluble protein GAPDH, calnexin (ER marker), and ZIKV protein prM were all present in the detergent-soluble fraction. In contrast, the majority of NS2B and NS5 were present in the detergent-resistant fraction of treatment (up to 1% Triton X-100) (Fig. 1B). For the detection of ZIKV dsRNA, which is supposed to be located inside the RCs, reverse transcription was performed with a forward primer to synthesize cDNA from the ZIKV negative-sense RNA ((-)RNA), followed by real-time PCR (RT-qPCR). The ZIKV dsRNA was primarily detected in the pellet samples after treatment with up to 1% IGEPAL CA-630 (Fig. 1C). These results suggest that the RCs may be present in the detergent-resistant fraction.

**Fig. 1.**
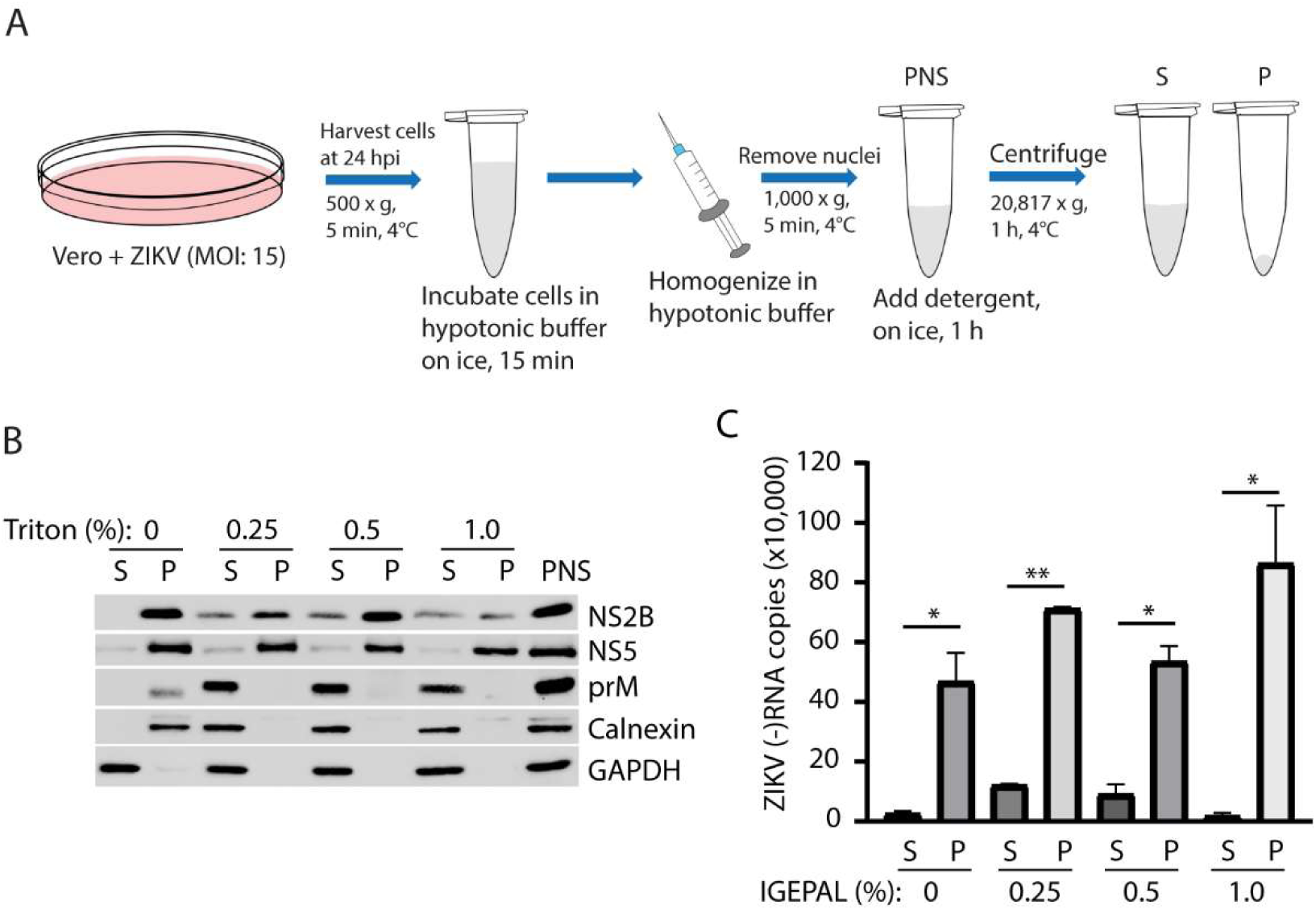
The detergent-resistant fraction of ZIKV-infected cells contains the replication complexes. A. Schematic illustration of the experimental procedure to isolate the post-nuclear supernatant (PNS) and treatment with detergent. Vero cells were inoculated with ZIKV PR strain at an MOI of 15, harvested at 24 hpi, and subjected to hypotonic buffer treatment for 15 min on ice. Cell homogenization was performed using a 1-ml syringe and a 25-gauge needle. The supernatant (S) and pellet (P) represent detergent-soluble and resistant fractions, respectively. B. Western blotting shows the detergent-resistant proteins in the pellet (P) and the detergent-soluble proteins in the supernatant (S). Triton X-100 was used at the indicated concentrations. Antibodies against ZIKV proteins NS2B, NS5, and prM, calnexin (ER marker), and GAPDH (soluble in cytosol) were used. C. Detection of the ZIKV (-)RNA in S and P portions. IGEPAL CA-630 was used at the indicated concentrations. Copies of total ZIKV (-)RNA in the respective fractions are shown. RNA levels between the S and P portions in each treatment are significantly different: * denotes *P* < 0.05, and ** denotes *P* < 0.01.

### ZIKV RCs are present in the top fraction of sucrose flotation

We isolated the RCs from ZIKV-infected Vero cells as described (16) with modifications (Fig. 2A). Briefly, approximately 6 × 10^6^ Vero cells were infected with ZIKV PR strain at an MOI of 15 and harvested 24 hpi. The cells were ruptured with detergent after incubation in hypotonic buffer, and the nuclei were removed by centrifugation. The cytoplasmic fraction containing membranous structures was mixed with 72% sucrose and placed at the bottom of the centrifugation tube, topped with 45% and 10% sucrose, which were subjected to flotation at 165,000 × g for 4 h. Fractions of 1 ml each were collected and used to analyze viral and cellular proteins by WB and to detect ZIKV dsRNA by RT-qPCR. WB results showed that ZIKV proteins NS2B, NS3, and NS5 were present in the top fraction, albeit at lower levels than those in fractions 3-5, in the sample from ZIKV-infected cells. In contrast, prM, calnexin (ER marker), succinate dehydrogenase subunit A (SDHA) (mitochondria marker), and GAPDH (cytoplasmic and soluble) were mainly present in fractions 3-5. Their presence in fraction 1 was below the detection limit (Fig. 2B). ZIKV proteins were absent in the mock-infected sample, and the distribution of calnexin, SDHA, and GAPDH was observed in fractions 3-5, similar to that in the infected cell sample. The ZIKV (-)RNA was mainly detected in the top fraction of the infected cell sample at the highest level (Fig. 2C). The results suggest that the RCs float to the top in the gradient centrifugation.

**Fig. 2.**
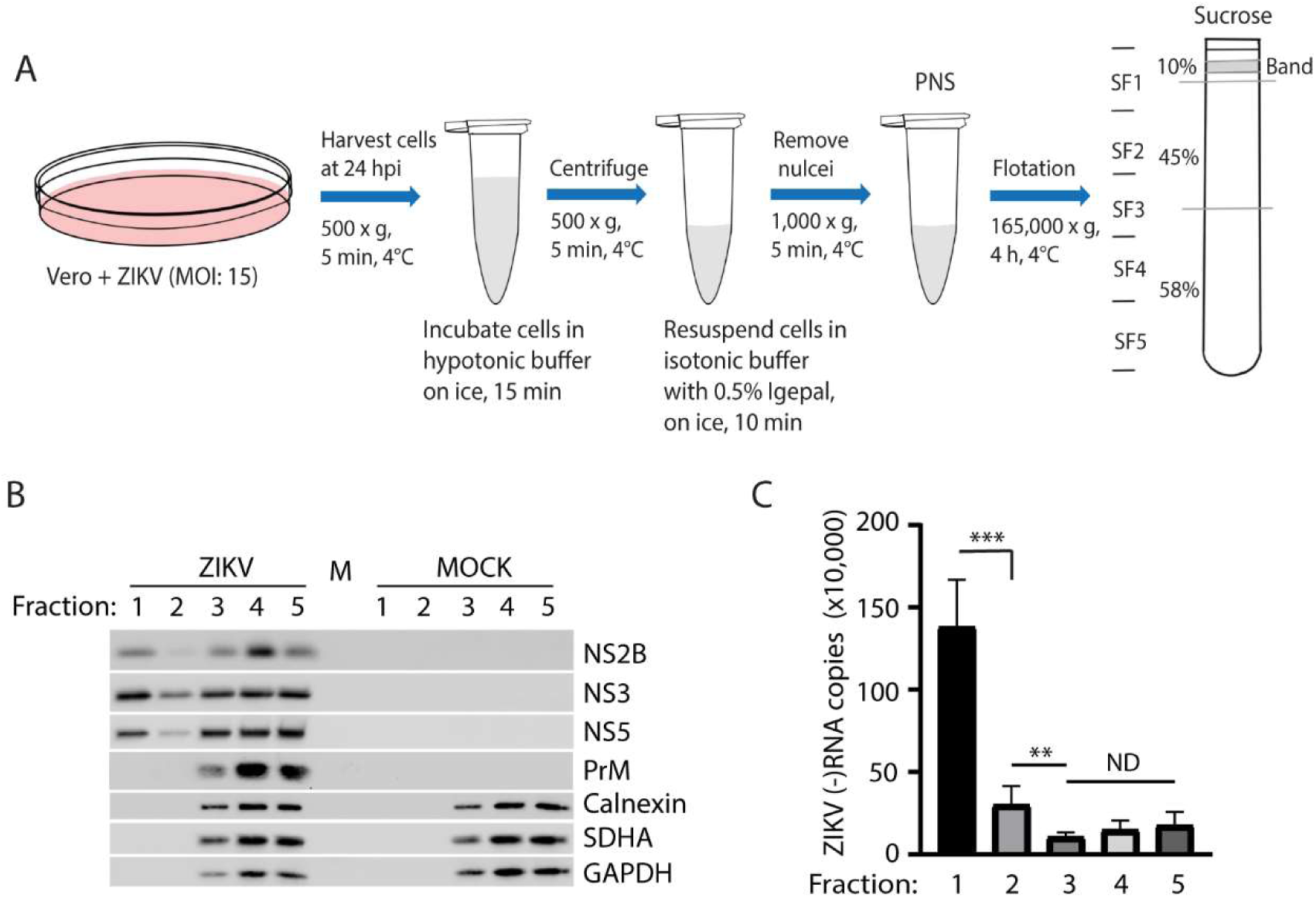
Isolation of the ZIKV RCs by sucrose flotation centrifugation. A. Schematic illustration of RC isolation from harvesting cells to sucrose density gradient centrifugation. PNS was mixed with sucrose to form a 58% solution and placed at the bottom of the centrifugation tube. Fractions SF1-SF5 were collected for WB and RNA isolation. B. Presence of ZIKV NS2B, NS3, and NS5 in the top fraction. Fractions SF1-SF5 were subjected to WB with antibodies against NS2B, NS3, NS5, prM, calnexin (ER marker), SDHA (mitochondrion marker), and GAPDH (soluble in the cytosol). M, molecular weight marker lane. C. Negative-sense ZIKV RNA in flotation fractions. Total copies of the ZIKV (-)RNA in each fraction are shown. Significant differences between fractions are indicated with *** for *P* < 0.001 and ** for *P* < 0.01. ND, no significant difference.

### The ZIKV RCs isolated by sucrose flotation are functional in viral RNA synthesis and remain active after cold storage

To determine if the isolated ZIKV RCs are functional in RNA synthesis, we conducted an *in vitro* RNA synthesis assay (16). The top fraction of the sucrose flotation was concentrated and used in the assay with the addition of a ribonucleotide solution set (NTP). The assay was performed at 30°C for 2 hours, followed by RNA isolation and RT-qPCR to determine the ZIKV (-)RNA yield. Results showed that, compared to the NTP-negative control, the addition of NTP led to a four-fold increase in ZIKV (-)RNA yield (Fig. 3A), demonstrating active RNA synthesis. To determine whether a stronger detergent, such as SDS, would destroy the RCs, we treated them with 1% SDS during the *in vitro* RNA synthesis assay. The addition of SDS abolishes the RCs’ synthesis of (-)RNA (Fig. 3B). To assess the stability of the isolated RCs, we store them at 4°C or −80°C for 24 hours before performing the *in vitro* RNA synthesis assay. The results showed that the RCs remained functional in ZIKV RNA synthesis after storage at 4°C and −80°C for 24 hours (Fig. 3C). Since the RCs isolated by sucrose flotation remained functional and stable after 24 h of cold storage, we proceeded with further purification.

**Fig. 3.**
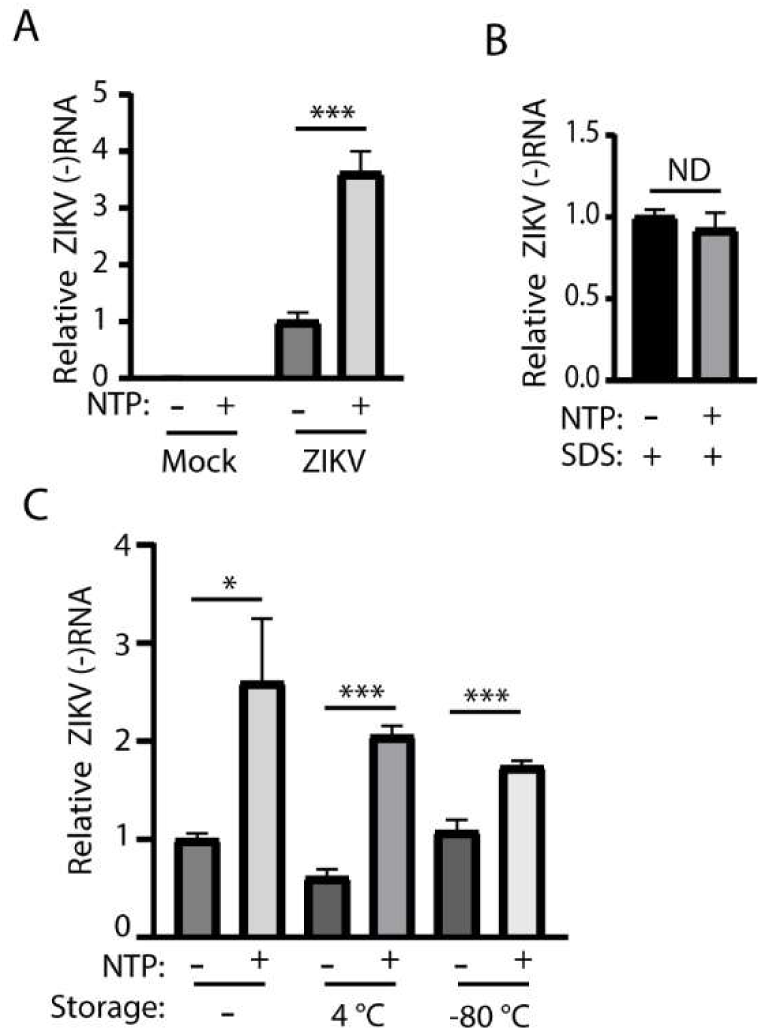
The RCs isolated from sucrose flotation are functional in RNA synthesis and remain active after cold storage. A. The top fraction is functional in RNA synthesis. The top fraction without NTP addition served as the control. Then, RNA was extracted for RT-qPCR. The representative of three experiments is shown. *** denotes *P* < 0.001. B. Treatment of the RCs with 1% SDS inactivates them in RNA synthesis. C. The RCs remain active in RNA synthesis after storage at 4°C or −80°C for 24 hours. * denotes *P* < 0.05.

### Purification of ZIKV RCs with iodixanol gradient flotation

To further purify the ZIKV RCs, we performed iodixanol gradient ultracentrifugation. The top fraction of sucrose flotation was mixed with an iodixanol solution to achieve a final concentration of 30%. The mixture was placed at the bottom of the centrifuge tubes, followed by consecutive layers of 25%, 20%, 15%, and 10% iodixanol, and centrifuged at 125,000 × g for 14 h (Fig. 4A). Fractions of 1 ml each were collected and used for the detection of ZIKV dsRNA by RT-qPCR. The results showed that fraction 3 contained the highest number of (-)RNA copies, followed by fractions 2 and 4 (Fig. 4B), indicating the predominant presence of RCs in these three fractions. The spread of the RCs across the three fractions was likely due to sampling, differences in size among individual RCs, or aggregation of multiple RCs. We then conducted an *in vitro* RNA synthesis assay with fraction 3. The freshly isolated RCs support new RNA synthesis, as indicated by a more than 4-fold increase upon the addition of NTPs compared to the control (Fig. 4C). These results demonstrate that we purified the RCs using iodixanol gradient flotation and that the newly isolated RCs are functional in RNA synthesis.

**Fig. 4.**
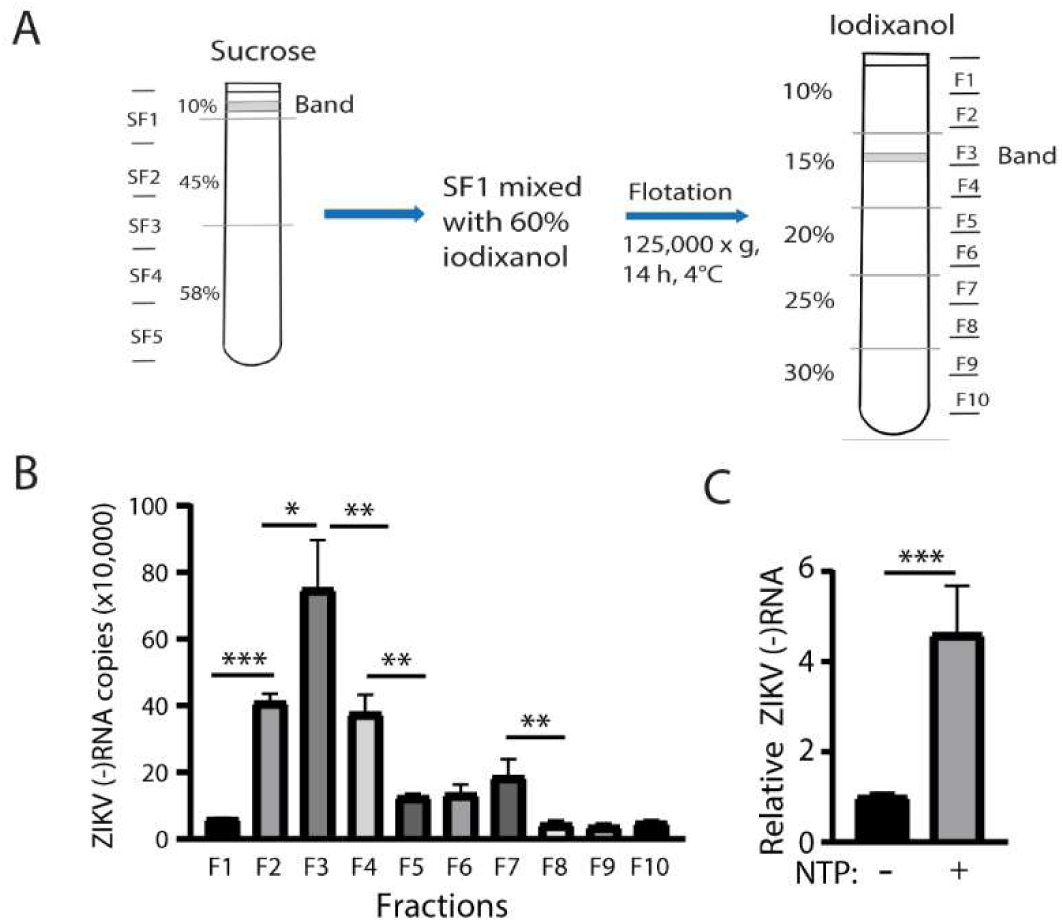
Purification of the ZIKV replication complexes by iodixanol gradient centrifugation. A. Schematic illustration of experimental procedures to purify RCs from the flotation of PNS in a sucrose gradient to the flotation of the sucrose top fraction in an iodixanol gradient. The membranous bands in the top fraction of sucrose and iodixanol gradients are indicated. The fractions collected are also indicated. B. Total ZIKV (-)RNA in the iodixanol fractions. Significant differences between fractions are indicated: * denotes *P* < 0.05; ** denotes P < 0.01; *** denotes *P* < 0.001. C. Purified RCs are functional with *in vitro* RNA synthesis activity.

### Mass spectrometry analysis of the purified RCs

After determining that the purified RCs remain functional in RNA synthesis, we wondered what host factors were present in the RCs. The total proteins in fraction 3 from iodixanol gradient flotation were then subjected to mass spectrometry (MS) analysis. A total of 769 proteins were identified in both mock and ZIKV-infected samples (Supplemental Table 1). Among them, 319 were shared between these two samples, 352 were unique to the ZIKV-infected sample, and 98 were unique to the mock-infected control (Fig. 5A). To explore the possible molecular mechanisms linked to the ZIKV replication, we performed a functional enrichment analysis of proteins from the mock (417 proteins) and ZIKV (671 proteins) samples using the PANTHER classification system against all Homo sapiens protein-coding genes using Fisher’s exact test (18).

**Fig. 5.**
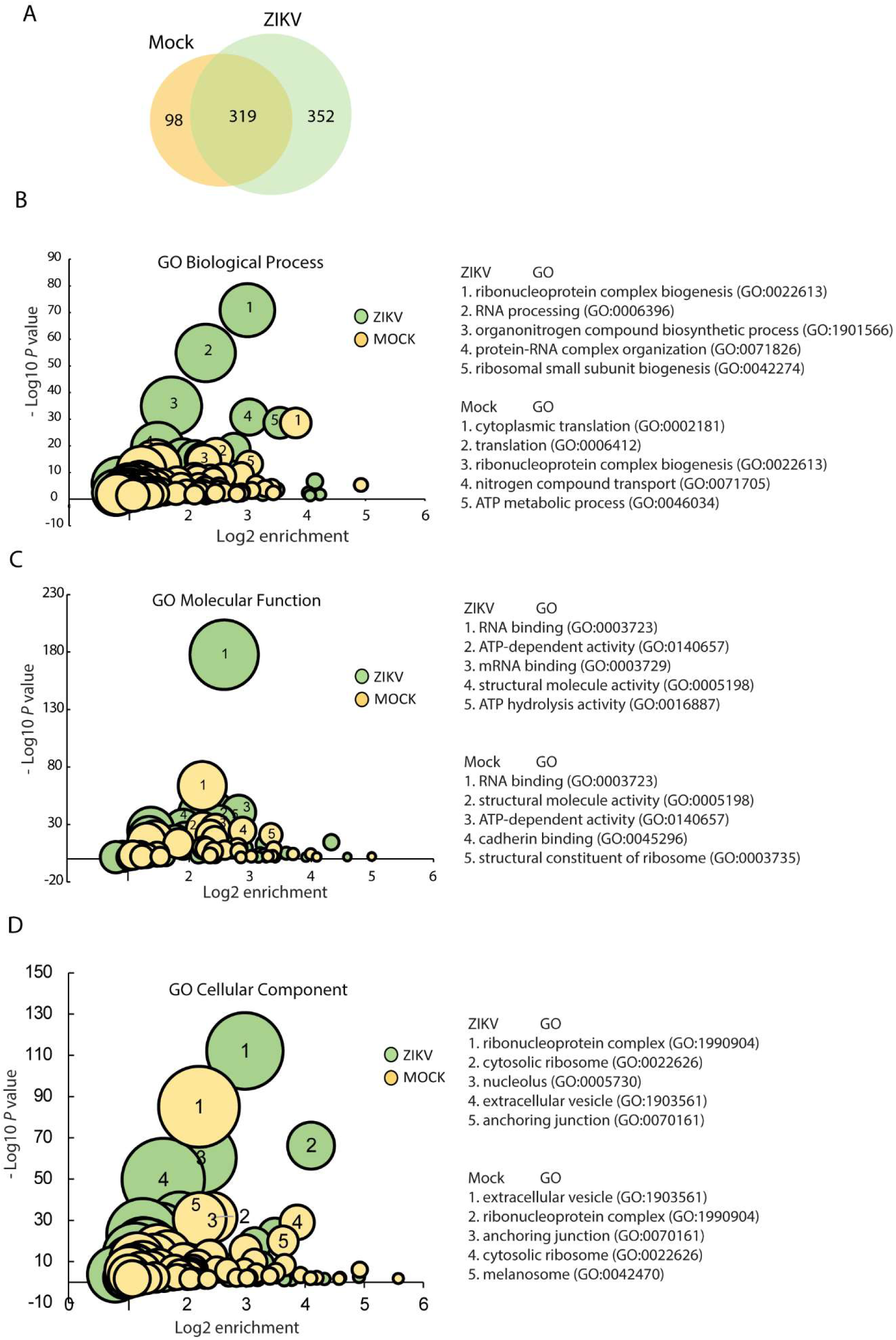
Gene Ontology (GO) enrichment analysis of total proteins identified in ZIKV or mock-infected samples. A. Venn diagram summarizing the number of proteins detected in fraction 3 of iodixanol flotation of mock-infected and ZIKV-infected samples. The overlapping area shows the proteins detected in both samples, whereas the non-overlapping area indicates the number of unique proteins in each respective sample. GO enrichment analysis of the biological process (B), molecular function (C), and cellular component (D) of the total proteins detected in each of the two samples. The horizontal axes are Log2 of enrichment, and the vertical axes are negative Log10 of *P*-value for statistical significance of observed enrichment. Bubble graphs show the number of proteins associated with a particular GO term (bubble size). The top 5 GO terms enriched in Mock and ZIKV samples are listed on the right of the bubble charts and indicated in the relevant bubbles.

The significantly enriched terms in biological processes (BP), cellular components (CC), and molecular functions (MF) are shown in bubble charts from gene ontology (GO) enrichment analysis, with the top 5 terms listed on the right (Fig. 5B-D). Regarding the enrichment of proteins by biological process, ZIKV RCs are highly enriched for processes that support an active response to infection, such as RNA processing and biosynthetic activities (Fig. 5B). In contrast, mock-infected cells focus on maintaining normal cellular functions, such as protein synthesis and energy metabolism. In molecular function, both samples involve RNA binding, but the ZIKV RCs sample showed a significant enrichment of RNA-binding proteins (331 total, Supplemental Table 2) (Fig. 5C). Regarding the cellular component, the enrichment of extracellular vesicles in both samples is anticipated due to the flotation-based isolation method, which naturally concentrates buoyant components, such as cholesterol-rich extracellular vesicles (Fig. 5D). Additionally, ZIKV RCs showed significant enrichment for other GO terms, such as ribonucleoprotein complexes and cytosolic ribosomes, indicating active RNA processing and protein synthesis, both of which are crucial for viral replication. Interestingly, several nuclear proteins were enriched in the ZIKV-infected samples. The analysis revealed that many proteins involved in RNA metabolism were potentially present in ZIKV-infected cells compared with the mock-infected control.

To gain deeper insight into proteomic remodeling induced by ZIKV infection, we performed a GO enrichment analysis using proteins detected exclusively in either the mock-infected control (98 proteins) or the ZIKV-infected sample (352 proteins) (Supplemental Table 3). Overall, GO term enrichment of proteins from the ZIKV RCs was significantly higher than that of proteins from mock-infected cells (Fig. 6A-C). In the mock-infected control, the enriched GO terms are consistent with the total protein enrichment analysis, in which the enriched proteins maintain regular metabolic and catabolic activities, extracellular communication, and transport (Supplemental Table 3). In the isolated ZIKV RCs, the proteins are highly enriched for RNA-processing and nuclear-associated components. Interestingly, half of the RNA-binding-enriched proteins are those exclusively identified in the ZIKV RCs. Regarding the enrichment of proteins based on molecular function, the top five GO terms in the sample of ZIKV RCs are RNA-binding, ATP-dependent activity, helicase or ATP hydrolysis, whereas the GO terms in the mock-infected control focus on antioxidant and oxidoreductase activity with a much smaller number of proteins (Fig. 6B), which suggests that ZIKV might recruit many cellular RNA-processing proteins for replication.

**Fig. 6.**
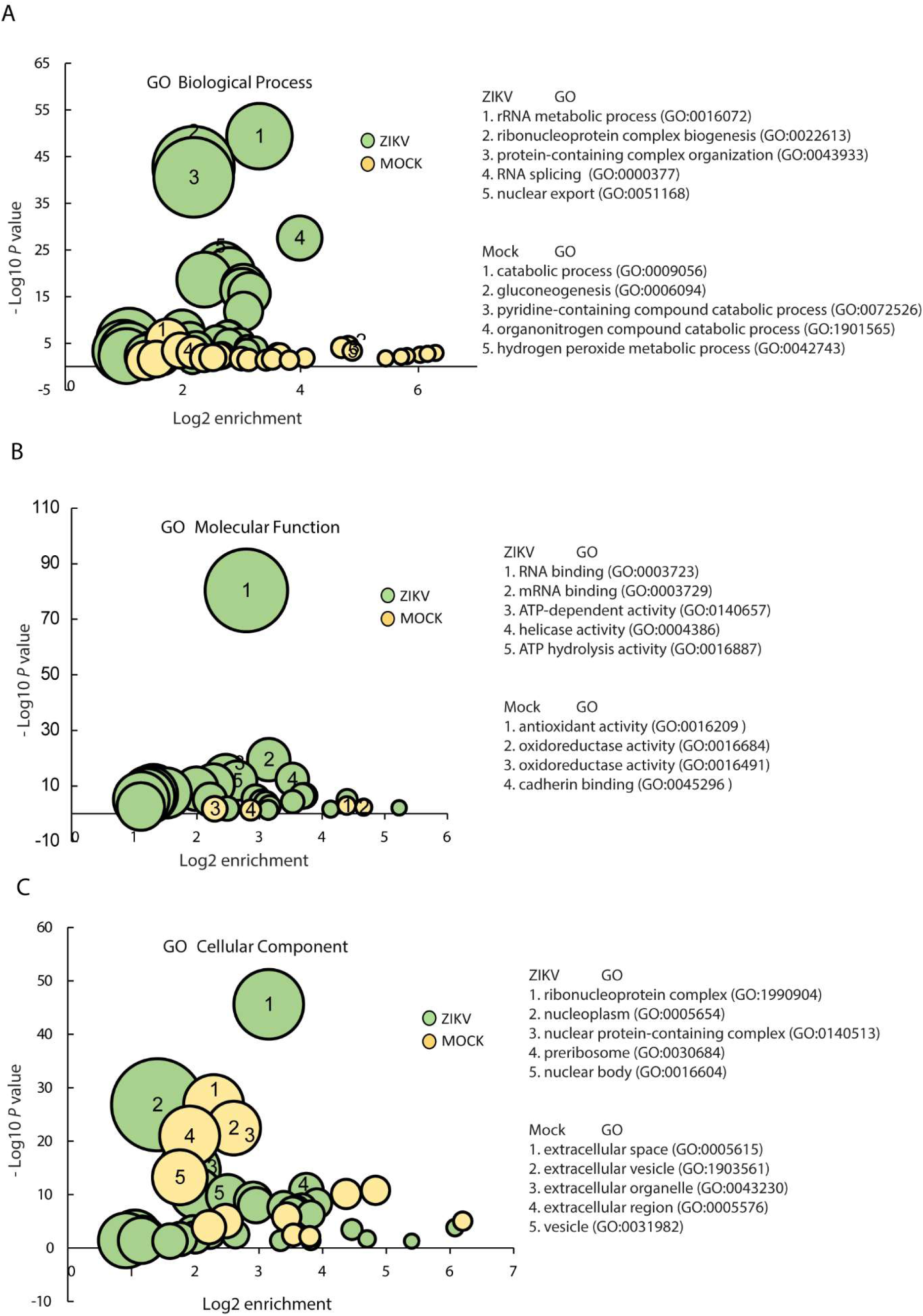
GO enrichment analysis of proteins specifically identified in ZIKV or Mock samples. GO enrichment analysis of the biological process (A), molecular function (B), and cellular component (C) is shown. The horizontal axes are the Log2 of enrichment, and the vertical axes are the negative Log10 of the *P*-value for the statistical significance of the observed enrichment. Bubble graphs show the number of proteins associated with a particular GO term (bubble size). The top 5 GO terms enriched in Mock and ZIKV samples are listed beside the bubble charts and indicated in the relevant bubbles.

Among the top hits exclusively in the ZIKV RCs sample, we noted a nuclear factor, replication factor C subunit 2 (RFC2), which is involved in the top molecular function GO terms, including RNA-binding, ATP-dependent activity, helicase, or ATP hydrolysis. We speculated that ZIKV might recruit this nuclear protein to the RCs to facilitate its replication. So, RFC2 was selected for further analysis.

### ZIKV infection elevates the RFC2 protein level and induces its relocation to the cytoplasm

Vero cells were infected with the ZIKV PR strain at an MOI of 1. WB results showed that the RFC2 level increases in the ZIKV-infected Vero cells at 24 hpi by 2.6-fold compared with the mock-infected cells (Fig. 7A). To determine if the ZIKV-induced RFC2 elevation is not limited to Vero cells, derived from African monkey kidney, we also used A549, human alveolar basal epithelial cells, for ZIKV infection. WB showed that ZIKV infection increased RFC2 levels in A549 cells by 5.9-fold compared with the control (Fig. 7B). These results demonstrate that ZIKV induces an elevation of RFC2 levels in infected cells. Since RFC2 is an accessory protein of DNA polymerase in the nucleus and was detected in isolated RCs, we wondered if the subcellular localization of RFC2 would be affected by ZIKV infection. An immunofluorescence assay (IFA) was performed on ZIKV-infected Vero cells using antibodies against RFC2 and ZIKV NS2B. The results showed that ZIKV infection induced the relocation of RFC2 to the cytoplasm (Fig. 7C). RFC2 appeared to colocalize with ZIKV NS2B, a viral protein apparently present in the RCs, with a Pearson’s correlation coefficient (PCC) of 0.74.

**Fig. 7.**
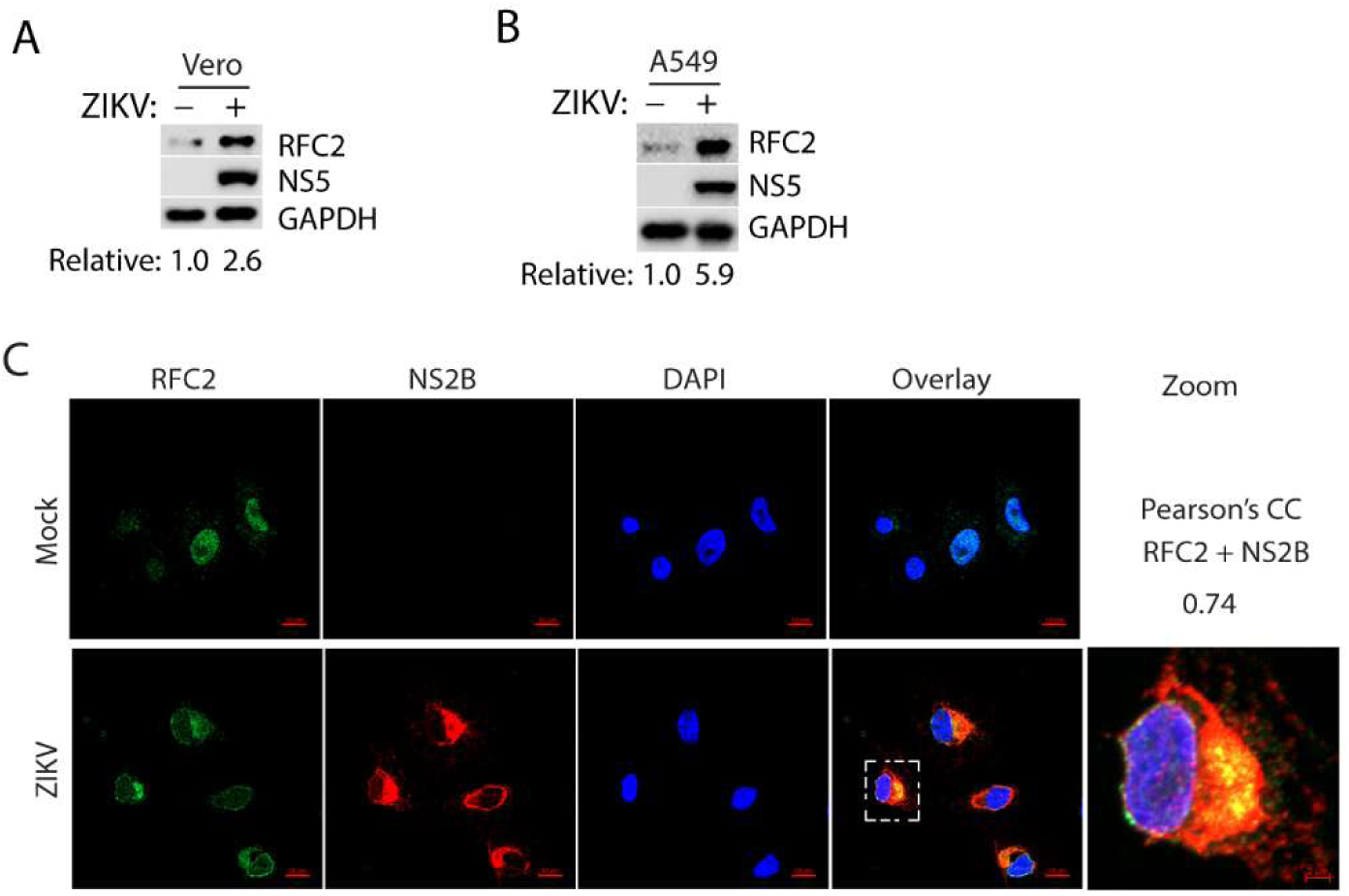
ZIKV infection induces the elevation of the RFC2 protein level and its relocation to the cytoplasm. A. RFC2 elevation in ZIKV-infected Vero cells. The cells were infected with ZIKV PR strain at an MOI of 1 and harvested 24 hpi for WB. B. RFC2 elevation in ZIKV-infected A549 cells. The cells were infected with ZIKV PR strain at an MOI of 0.1 and harvested 24 hpi for WB. Densitometric analysis was performed, and relative RFC2 levels, normalized to GAPDH, are shown below the images. C. ZIKV infection induces RFC2 relocation to the cytoplasm in Vero cells. The cells were infected with the ZIKV PR strain at an MOI of 1 and fixed 24 hpi for IFA. Mock-infected cells were included as a control. Overlay of RFC2, NS2B, and DAPI is shown. The scale bars in the lower right of each image denote 10 μm. One cell was cropped to show the zoom-in. The scale bar in the lower right of the zoomed-in image denotes 2 μm. PCC is shown above the image.

The results above prompted us to assess the effect of RFC2 on ZIKV replication. We designed two small hairpin RNAs (shRNAs) against RFC2 and cloned them into a retroviral expression system. RNAi-mediated silencing of RFC2 was performed in SVG astrocytes, a physiologically relevant cell type for ZIKV infection. WB results showed that, compared with the control shRNA, shRFC2-1 and shRFC2-2 reduced RFC2 levels to 0.12- and 0.05-fold, respectively, in SVG cells (Fig. 8A). A cell viability assay was conducted to ensure that RFC2 depletion had minimal effects on cell growth. Indeed, the SVG cells with RFC2 depletion proliferated similarly to cells treated with the control shRNA (Fig. 8B). We then infected the RFC2-depleted cells with the ZIKV PR strain and harvested them 24 hpi for WB and RT-qPCR. The WB result showed that ZIKV NS5 protein levels in the RFC2-depleted cells were significantly lower than that in the control cells, with reductions to 0.45- and 0.19-fold for the cells treated with shRFC2-1 and shRFC2-2, respectively (Fig. 8C). RT-qPCR results showed that the cells with RFC2 depletion had significantly lower ZIKV RNA levels than the control cells, with a significant reduction to 13.1% and 5.6% for the cells treated with shRFC2-1 and shRFC2-2, respectively (Fig. 8D). These results demonstrate that RFC2 depletion impairs ZIKV replication, suggesting RFC2 is possibly a proviral factor.

**Fig. 8.**
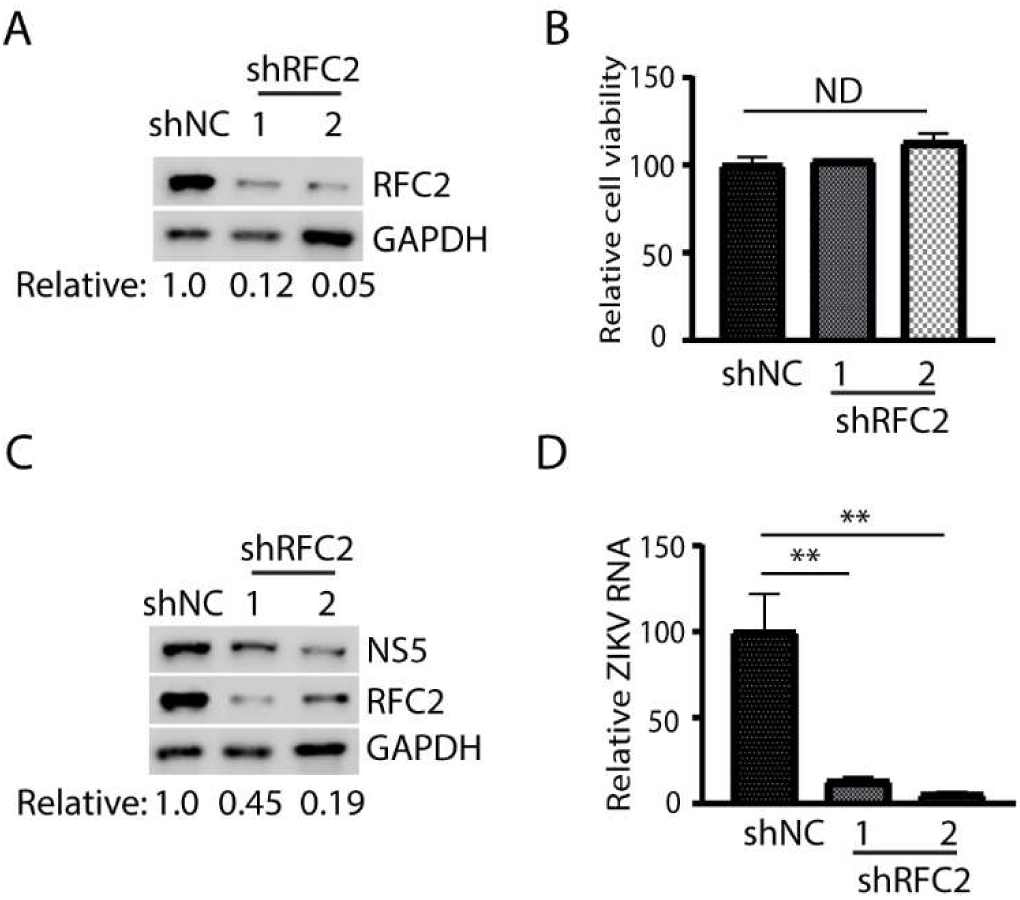
RFC2 depletion impairs ZIKV replication in SVG cells. A. RNAi-mediated RFC2 depletion in SVG cells. Relative RFC2 levels, normalized to GAPDH, are shown below the images. shNC: control shRNA. shRFC2, shRNA against RFC2. B. Relative cell viability of SVG cells. The relative cell viability percentages compared with shNC-treated cells are shown. ND, no significant difference. C. RFC2 depletion reduces ZIKV NS5 protein levels. SVG cells were inoculated with ZIKV PR strain at an MOI of 1 and harvested for WB at 24 hpi. Relative NS5 levels, after normalization to GAPDH, are shown below the images. D. Depletion of RFC2 reduces ZIKV RNA levels. SVG cells were transduced with a recombinant retrovirus containing shRNA three times over two days per passage. The cells were then inoculated with ZIKV at an MOI of 5 and harvested 24 hpi for RNA isolation. The relative percentages of ZIKV RNA, compared with those in shNC-treated cells, are shown. Significant differences between groups are indicated by ** (P < 0.01).

## DISCUSSION

In this study, we isolated and purified ZIKV RCs, as evidenced by the presence of viral proteins and negative-sense RNA in the same flotation fraction. In addition, purified RCs retain functions in RNA synthesis and stability at low temperatures. Proteomic analysis identified potential host factors in RCs, contributing to our understanding of how ZIKV manipulates host cellular machinery for replication and revealing potential targets for antiviral research.

Previous studies used transmission electron microscopy to observe the morphology of the ZIKV RCs or immunofluorescence assays to detect viral double-stranded RNA (dsRNA), thereby indirectly indicating its presence (5, 19–22). These methods provide some insights, but direct isolation is needed to study the molecular mechanism of ZIKV RCs. In the first step, we must obtain the cell lysate containing the RCs. The mechanical technique of disrupting cells with a needle and syringe or a Dounce homogenizer is often employed, but it presents several challenges (16, 23). This process is time-consuming and can result in sample loss.

Additionally, the strength and speed used during this procedure can significantly affect the final results, potentially introducing variability or compromising the integrity of the isolated components. It is known that, like other (+)RNA viruses, ZIKV requires fatty acids, cholesterol, glycerophospholipids, phospholipids, and sphingolipids (mainly ceramides) to form the RCs (9, 10, 24). Taking advantage of the relatively detergent-resistant nature of cholesterol-rich lipids, we decided to lyse cells with the nonionic detergent Triton X-100, which gently solubilizes membrane lipid components. This treatment helps release the RCs while preserving their structure and associated proteins. This strategy was validated by detecting negative-sense viral RNA and viral proteins (NS2B, NS5) in the detergent-resistant fraction of the cell lysate (Fig. 1B-C). Like the soluble protein GAPDH, some membrane-bound proteins not involved in the RCs, such as calnexin in the ER and ZIKV prM, were released from the membrane by the detergent treatment (Fig. 1B). Although ZIKV prM mediates viral entry and assembly by binding with cholesterol (25), our results suggest their interaction is less stable or transient. So, we used a nonionic detergent for the RCs isolation. In addition, cells were incubated in a hypotonic buffer before lysis, thereby facilitating detergent-containing isotonic lysis. The next step was sucrose flotation centrifugation to enrich detergent-resistant membranous structures in the top fraction while retaining soluble components in the bottom fractions. Further purification of the RCs was achieved by flotation of the top sucrose fraction in an iodixanol density gradient, yielding cleaner, enriched RCs. Our purified RCs were functional in ZIKV RNA synthesis.

The viral proteins NS2B, NS3, and NS5 were detected in the top fraction of the sucrose flotation assay (Fig. 2). However, they were also detected in the bottom fractions. The possible reason is that these proteins are not present solely in the RCs. ZIKV NS5 predominantly accumulates in the nucleus, although ZIKV replication occurs in the cytoplasm (26). NS5 is indispensable for ZIKV RNA synthesis because it contains an N-terminal methyltransferase (MTase) domain responsible for viral RNA capping and a C-terminal RdRp domain for viral RNA synthesis (13, 27). The nuclear accumulation protects NS5 from cytoplasmic degradation (26). Detergent-lysed cells may release NS5 from the nucleus, which cannot float as the NS5 is non-membrane-enclosed. NS2B is a transmembrane protein that recruits NS3 to RCs and functions as a cofactor for the protease activity of NS3 (28). Without the transmembrane domain and interaction with NS2B, NS3 localizes to the mitochondria. Understandably, not all the NS3 proteins are in RCs. When independent of the NS2B-NS3 complex, NS2B has additional functions, including mediating the interaction between protein phosphatase 1α (PP1α) and eukaryotic initiation factor 2α (eIF2α), promoting eIF2α dephosphorylation, and inhibiting stress granule formation (29). The Dengue virus NS2B was found to interact with MAVS and IKKε to impair RIG-I-directed antiviral response (30). NS2B may be free from RCs under those interactions in the cells, resulting in solubilized NS2B in the bottom fractions.

Our purification of the RCs resulted in a ZIKV replication-competent fraction enriched for the negative-sense RNA, as shown in the *in vitro* RNA replication assay (Fig. 4C). Like other viruses, ZIKV is an obligate intracellular agent that depends on host-cell metabolites and machinery to produce infectious progeny virions. To determine the RC components, we performed mass spectrometry analysis of the purified RCs. It is not surprising that ZIKV hijacked many host proteins for its RCs (Fig. 5A and Supplemental Table 1). Given that these samples were subjected to detergent treatment and gradient flotation, the detection of membrane-bound components in the floated fraction from the mock-infected control is expected. It indicates the presence of cholesterol-rich structures, implying that ZIKV infection may alter the abundance or utilization of pre-existing components.

Among the proteins detected in the RCs, many are nuclear, suggesting that the virus recruits or redistributes them to cytoplasmic replication complexes. GO enrichment analysis of proteins exclusively in the ZIKV RCs or control confirmed that those nuclear proteins are almost exclusively in the ZIKV RCs (Fig. 6). This indicates that ZIKV may hijack some of the nuclear functions, like RNA processing (e.g., ribonucleoprotein complex biogenesis and RNA splicing), or disrupt nuclear-cytoplasmic transport to support its replication (nuclear export). It was reported that ZIKV infection relocates the nuclear protein high-mobility group box 1 (HMGB1) into the cytosol to facilitate its replication (31). HMGB1 was also detected in our isolated RCs, consistent with the previous study.

Previous research identified some host factors required for the ZIKV RCs (20, 32, 33). The proteins identified in the RCs in our MS analysis are consistent with those identified in previous publications. For example, receptor for activated protein C kinase 1 (RACK1) (32), vimentin (33), and insulin-like growth factor 2 mRNA-binding protein 2 (IGF2BP2) (20) were also identified in the ZIKV RCs sample at a much higher abundance than in the control. ER membrane protein complex (EMC) is identified as a critical host factor for DENV and ZIKV replication, and NS4A/NS4B requires the EMC for their efficient biogenesis (34). We identified four EMC proteins (EMC1-3 and EMC7) in the ZIKV RC sample, all of which showed a significant elevation in abundance compared to the mock-infected control. EMC7 was exclusively found in the RCs. Fragile X mental retardation syndrome-related protein 2 (FXR2) is an RNA-binding protein and was also found exclusively in the ZIKV RCs. Previous research confirmed the colocalization of FXR2 with ZIKV NS4B and showed that FXR2 knockdown impairs ZIKV replication (35). Thus, being consistent with these earlier works supports our RC isolation, which appears reliable for its intended application.

RFC2 is one of five subunits (RFC1–RFC5) of the RFC complex, a DNA polymerase accessory factor essential for processive DNA elongation (30–32). All subunits contain conserved ATP-binding and hydrolysis domains, and homologous proteins exist across bacteria and phages (33). RFC2 interacts with RFC1 and RFC5 within the complex and with DNA polymerases δ and ε during replication (31, 34). The RFC complex loads the DNA polymerase processivity factor, proliferating cell nuclear antigen (PCNA), onto DNA via ATP hydrolysis, and RFC2 is essential for RFC ATPase activity and PCNA loading onto chromatin (32). In addition to the canonical RFC1–RFC complex, RFC-like complexes (RLCs) composed of RFC2–5 and alternative large subunits (CTF18, RAD17, ATAD5/Elg1) function as PCNA loaders during DNA replication and DNA damage responses or PCNA removers after DNA synthesis (35–38). Thus, RFC2 participates in DNA replication, DNA repair, and cell-cycle checkpoint signaling (31, 39). Consistent with these roles, depletion of RFC2 causes chromosomal segregation defects and reduced cell numbers in neonatal rat cardiac myocytes (40).

As an accessory factor for DNA polymerases δ and ε, RFC is recruited by DNA viruses that replicate in the nucleus. Simian virus 40 (SV40) requires RFC during DNA replication elongation (41). The adeno-associated virus (AAV) depends on RFC for DNA replication (42). Kaposi’s sarcoma-associated herpesvirus (KSHV) recruits RFC via LANA for DNA replication and episome maintenance (43). The hepatitis B virus (HBV) uses RFC and other host factors for plus-strand repair during cccDNA formation (44). In contrast, most RNA viruses, including flaviviruses, replicate in the cytoplasm. This is the first report to demonstrate the involvement of an RFC complex member in RNA virus replication.

In conclusion, we isolated ZIKV RCs and characterized their protein composition, providing crucial insights into the host proteins hijacked by ZIKV for replication. Among the host factors is RFC2, which appears to be proviral as its depletion impairs ZIKV replication. These results enhance our understanding of virus-host interactions and provide valuable insights for future studies exploring viral mechanisms and developing antiviral strategies.

## MATERIALS AND METHODS

### Cells, viruses, and chemicals

Vero (ATCC CCL81) and A549 (ATCC CCL185) cells were maintained in Dulbecco’s Modified Eagle Medium (DMEM) supplemented with 10% fetal bovine serum (FBS). SVG (ATCC CRL8621) cells were maintained in Minimum Essential Medium (MEM) supplemented with 10% FBS. Cells were grown at 37°C with 5% CO_2_. ZIKV strain PRVABC59 (ATCC VR-1843) was used in this study. Nonionic detergent Triton X-100 (Sigma-Aldrich, St. Louis, MO) and IGEPAL CA-630 (Sigma-Aldrich) were used to lyse cells. Iodixanol solution (60%, Sigma-Aldrich) was used for gradient centrifugation in the purification of replication complexes.

### Detergent Resistance Assay

About 3 × 10^6^ Vero cells were infected with ZIKV at an MOI of 15. The cells were harvested at 24 hpi using a cell scraper, rinsed with cold PBS (pH 7.4), and incubated in 1 mL of hypotonic buffer (10 mM Tris-HCl (pH 7.5), 10 mM KCl, 5 mM MgCl_2_) for 15 min on ice. Then, the cells were pelleted by centrifugation and resuspended in 500 L of hypotonic buffer containing a protease inhibitor cocktail (Sigma-Aldrich) and RiboLock RNase inhibitor (Thermo Fisher Scientific, Waltham, MA). Cells were disrupted using a 25-gauge needle and syringe by 30 up-and-down strokes. The unlysed cells and nuclei were removed by centrifugation at 1000 × g for 5 min at 4°C, yielding the PNS. To determine the detergent sensitivity, the PNS was aliquoted and incubated with different concentrations of Triton X-100 or IGEPAL CA-630 (0, 0.25%, 0.5%, 1%) (v/v) on ice for 1 hour. After incubation, the soluble and insoluble parts were separated by centrifugation at 20,817 × g for 1 hour at 4 °C, yielding the supernatant (S) and pellet (P), which were used for total RNA extraction and WB.

### Isolation of the ZIKV replication complexes

About 6×10^6^ Vero cells in two 10-cm dishes were infected with ZIKV at an MOI of 15. Mock-infected Vero cells served as a control. At 24 hpi, the cells were harvested by scraping in cold PBS and rinsed. Cells were incubated in hypotonic buffer on ice for 15 min, then centrifuged. The cells were incubated with 500 μL isotonic buffer (50 mM Tris-HCl (pH 7.5), 150 mM NaCl, 2 mM EGTA, 50 mM MgCl_2,_ 1 mM sodium Vanadate, 10% Glycerol) supplemented with protease inhibitor cocktail and RNase inhibitors and 0.5% IGEPAL CA-630. After incubation on ice for 10 min, unlysed cells and nuclei were removed by centrifugation at 1000 × g for 5 min at 4°C. The PNS (500 μL) was mixed with 2 mL 72% sucrose solution (diluted in 50 mM Tris-HCl (pH 7.5), 25 mM KCl, 5 mM MgCl_2_), overlaid with 2 mL 45% sucrose dilution, and topped with 0.5 mL 10% sucrose solution, followed by centrifugation in a SW55Ti rotor at 165,000 × g for 4 hours at 4°C. After the flotation centrifugation, the sucrose gradients were fractionated at 1 mL per fraction for further analysis.

### Purification of the ZIKV replication complexes

For further purification, the top fraction from the sucrose flotation was collected and purified by iodixanol gradient flotation. One mL top sucrose fraction was mixed with 1 mL 60% iodixanol (diluted in 10 mM Tris-HCl (pH 7.5), 10 mM KCl, 5 mM MgCl2, 250 mM sucrose), overlaid consecutively with 2 mL of 25%, 20%, 15%, and 10% iodixanol solution, followed by ultracentrifugation in a SW41Ti rotor at 125,000 ×g for 14 hours at 4°C. The iodixanol gradients were fractionated at 1 mL per fraction for further analysis. The fraction containing RCs and the same fraction from the mock-infected control were concentrated with Amicon^®^ ultra-0.5 centrifugal filter (100 kDa MWCO, MilliporeSigma, Burlington, MA) and subjected to mass spectrometry analysis at the Taplin Mass Spectrometry Facility, Harvard Medical School (Boston, MA). Only proteins identified by at least two unique peptides were retained for downstream proteomic analysis.

### Stability test of the replication complexes

To assess the stability of the purified RCs, aliquots were incubated at 4°C or −80°C for 24 hours, then subjected to an *in vitro* RNA synthesis assay, total RNA extraction, and RT-qPCR.

### *In vitro* RNA synthesis assay

The RNA synthesis assay was performed as previously detailed, with some modifications (36). After collecting the fractions containing the replication complex, each fraction was concentrated using an Amicon^®^ ultra-0.5 centrifugal filter (100 kDa MWCO). A standard 50 μL reaction mixture containing 50 mM HEPES (pH 7.5), 10 mM KCl, 10 mM MgCl_2_, 100 U RNase inhibitor, 2 mM DTT, 10 mg/mL Actinomycin D, 0.5 mM ATP, 0.5 mM CTP, 0.5 mM GTP, 0.5 mM UTP, and the concentrated RCs to make it up to 50 μL. The reaction mixtures were incubated at 30°C for 2 h, and RNA was extracted by phenol-chloroform extraction followed by ethanol precipitation. To test the sensitivity of the isolated RCs to sodium dodecyl sulfate (SDS), 1% of SDS (v/v) was added to the 50 μL reaction mixture, which was incubated at 30°C for 2 h, followed by RNA extraction.

### RNA isolation and real-time PCR

RNA from the supernatant, pellet, and flotation fractions, was extracted with TRIzol™ LS Reagent (Thermo Fisher Scientific) following the manufacturer’s instructions. For each flotation fraction, 250 μL of each sample was used for RNA extraction. RNA from ZIKV-infected cells was extracted with PureLink RNA Mini Kit (Thermo Fisher Scientific). Reverse transcription and real-time PCR (RT-qPCR) were performed as described previously (37, 38). In addition, a standard curve was generated using plasmids encoding ZIKV NS5 (37) to quantify viral RNA copy number. The real-time PCR primers used for ZIKV are listed in Table 1.

**Table 1.**
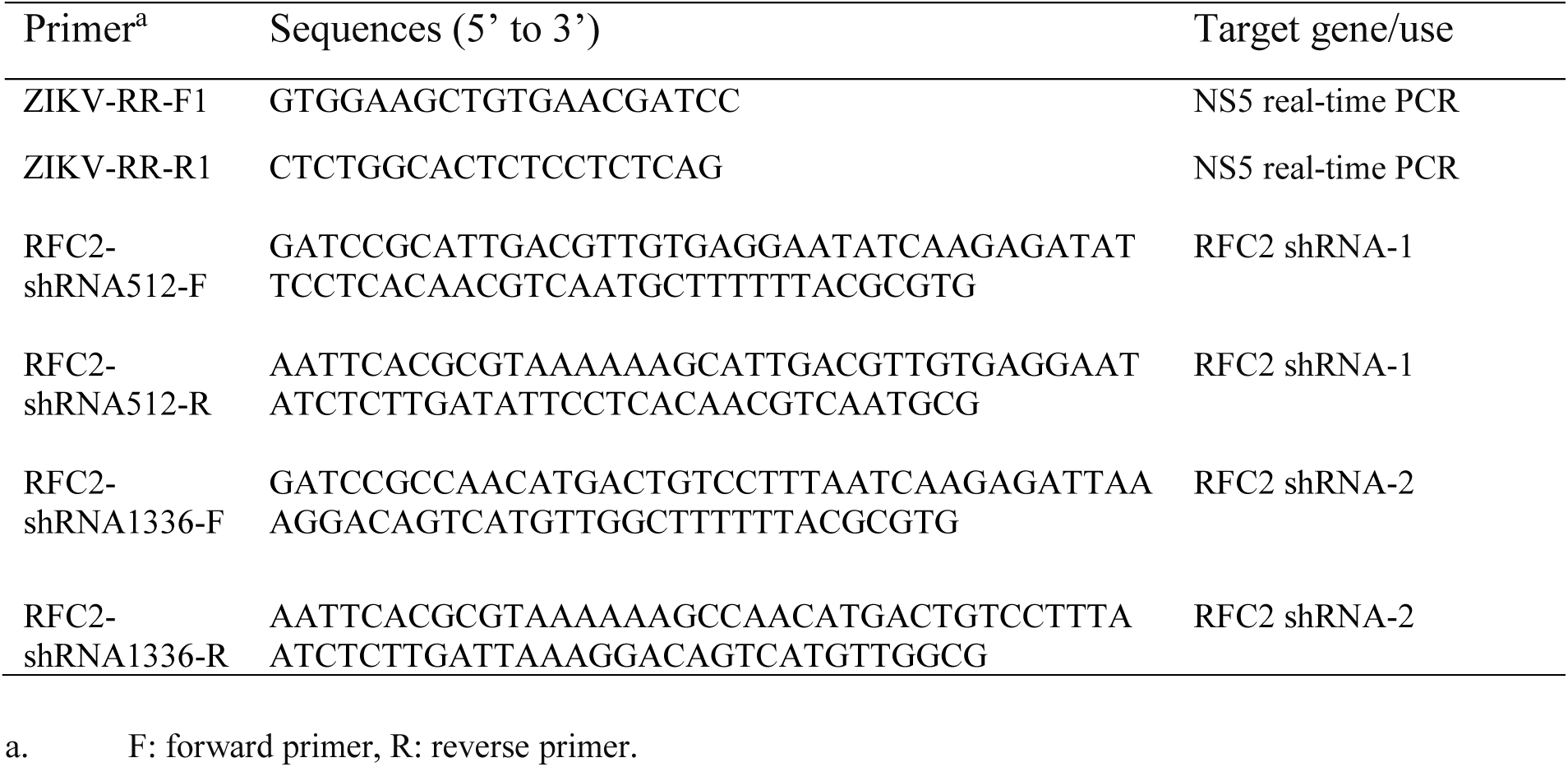
Primers used in this study.

### Western blotting (WB)

Cell lysates or other samples were subjected to SDS-PAGE and WB as described previously (37–39). The primary antibodies against ZIKV NS2B (GeneTex, Inc., Irvine, CA), NS3 (GeneTex), NS5 (GeneTex), prM (GeneTex), Calnexin (Santa Cruz Biotechnology, Inc., Dallas, TX), SDHA (Santa Cruz), GAPDH (Santa Cruz), and RFC2 (Proteintech Group, Inc., Rosemont, IL) were used in this study. The secondary antibodies conjugated with horseradish peroxidase include goat anti-mouse or anti-rabbit IgG (Rockland Immunologicals, Inc., Limerick, PA). The chemiluminescence substrate was used to reveal specific reactions, and the resulting signals were acquired within the linear range of digital intensity without pixel saturation. Densitometric analysis of the WB images was performed using the QuantityOne Program, version 4.6 (Bio-Rad Laboratories, Hercules, CA).

### Proteomics analysis

Gene Ontology (GO) enrichment of biological process, molecular function, and cellular component for all detected proteins in the mock and ZIKV-infected samples was analyzed using the PANTHER classification system web tool (https://pantherdb.org/) (18) against all Homo sapiens protein-coding genes using Fisher’s exact test and Bonferroni correction for multiple testing. *P*-values < 0.05 indicate statistically significant enrichment in specific GO terms. To reduce redundancy in GO terms, PANTHER GO enrichment results were imported into the REVIGO database (http://revigo.irb.hr/) (40).

### Plasmids

Oligos for shRNA (Table 1) were annealed and ligated into pSIREN-RetroQ-ZsGreen as instructed (Takara Bio USA, San Jose, CA) and previously described (41). The in-house-prepared plasmids were subjected to DNA sequencing by Plasmidsaurus using Oxford Nanopore Technology, with custom analysis and annotation.

### Immunofluorescence assay (IFA)

IFA was performed as previously reported (38). Primary antibodies against RFC2 (Proteintech) and ZIKV NS2B (GeneTex) and fluorescein-conjugated secondary antibodies, including goat anti-rabbit IgG (H&L) Dylight 549 and goat anti-mouse IgG (H&L) Dylight 488 (Rockland), were used. A Zeiss LSM800 confocal microscope and Zen software were used for imaging and evaluating the degree of colocalization of two fluorophores, as indicated by Pearson’s correlation coefficient (PCC): a value of 1 indicates perfect correlation, 0 for no correlation, and −1 for an ideal anti-correlation.

### Statistical analysis

Differences between the two groups were assessed by using Student’s t-test. A two-tailed *P* value of 0.05 was considered significant.

## Acknowledgment

This study was partially funded by a seed grant from the University of Maryland, College Park, MD. We thank Maria Salvato for her discussion and review of this manuscript. PC and BTS were supported by the Basil & Anne Hatziolos Scholarship Fund and the Barbara Holton Terwilliger Scholarship Fund for Veterinary Medical Research.

## Author Contributions

PC and YZ contributed to the study design and manuscript draft. PC, BTS, and YZ were involved in data acquisition, analysis, and manuscript modification. All authors reviewed the manuscript.

## Data Availability

Data are presented in this manuscript and the supplemental materials.

